# The TAT Protein Transduction Domain as an Intra-articular Drug-Delivery Technology

**DOI:** 10.1101/2020.04.28.066902

**Authors:** Sarah E. Mailhiot, Matthew A. Thompson, Akiko E. Eguchi, Sabrina E. Dinkel, Martin K. Lotz, Steven F. Dowdy, Ronald K. June

## Abstract

**Objective:** Intra-articular drug delivery holds great promise for the treatment of joint diseases such as osteoarthritis. The objective of this study was to evaluate the TAT peptide transduction domain (TAT-PTD) as a potential intra-articular drug delivery technology for synovial joints.

**Design:** Experiments examined the ability of TAT conjugates to associate with primary chondrocytes and alter cellular function both *in vitro* and *in vivo*. Further experiments examined the ability of the TAT-PTD to bind to human osteoarthritic cartilage.

**Results:** The results show that the TAT-PTD associates with chondrocytes, is capable of delivering siRNA for chondrocyte gene knockdown, and that the recombinant enzyme TAT-Cre is capable of inducing *in vivo* genetic recombination within the knee joint in a reporter mouse model. Lastly, binding studies show that osteoarthritic cartilage preferentially uptakes the TAT-PTD from solution.

**Conclusions:** The results suggest that the TAT-PTD is a promising delivery strategy for intra-articular therapeutics.

## Introduction

Intra-articular drug delivery is important for targeting therapeutic compounds to the diverse cells and tissues involved in arthritis pathogenesis (*i*.*e*. cartilage, bone, and synovium). Joint vasculature includes large pores (∼1-3 μm) in synovial blood vessels leading to fenestrated capillaries^4,17.^ These, as well as lymphatic drainage^9,25^, create challenges for drug residence times. Measurements show that most unmodified molecules do not possess sufficient joint residence time for effective pharmacologic intervention ^8,15,21,22,32,^ demonstrating a need for new delivery strategies.

Previous studies have explored synovial joint drug delivery using nanoparticles, microspheres, and phase-transitioning polymers ^6,13,19,24,28.^ Diclofenac sodium delivered by poly(lactic-co-glycolic acid) (PLGA) microspheres to antigen-induced chronic arthritic rabbits formulated for experimentally optimized conditions for extended release found no significant difference between PLGA treatment groups and control ^28^. Zilretta is an FDA-approved corticosteroid formulation for OA that achieves extended release through PLGA (poly lactic-co-glycolic acid) microspheres ^18,20^. Elastin-like polypeptides (ELPs), a type of phase-transitioning polymers set to aggregate at 37? following intra-articular injection, were found to have increased residency time for protein drug delivery during biodistribution tests in rats versus a soluble ELP control ^6^. These are exciting technologies show partial success in animal models.

The TAT peptide transduction domain (TAT-PTD) is a small, positively charged peptide that is electrostatically attracted to the negatively charged cell surface ^30^ and induces uptake in all cell types by binding to an as yet unidentified surface protein(s) ^12^. PTDs have enabled delivery of small and large (∼150 kDa) macromolecules into cells and have been tested in >2000 patients in over 25 clinical trials ^29^ and resulted in 2 statistically-significant Phase II trials, one published ^11^. Previous studies suggest that TAT-PTD transduction may be effective for modulating joint biology ^16,23,^ and PTDs remain to be systematically evaluated for joint disease applications.

Therefore, the objectives of this study were to first determine if the TAT-PTD could transduce macromolecules into chondrocytes, and second, to assess the ability of the TAT-PTD to bind to cartilage. Toward these goals, this study applied both TAT-conjugated peptides and proteins in *in vitro* and *in vivo* models. The result show that the TAT-PTD is capable of inducing transduction and binding to the osteoarthritis (OA) cartilage matrix. Future studies may build on these results to apply the TAT-PTD toward drug delivery in OA.

## Materials and Methods

### Primary Human Chondrocytes

This study utilized primary chondrocytes obtained with IRB approval and harvested from human knee joint femoral and tibial cartilage with Outerbridge grades 1-4. Sources of joint tissue included both healthy tissues obtained from tissue banks from patients without history of arthritis or any macroscopic signs of joint degeneration and osteoarthritic patients undergoing joint replacement surgery. Primary human chondrocytes (mean age: 64.6 years, age range: 36-94, 73% male and 27% female) were harvested from the joint by dissecting the cartilage as described ^1^, and cultured in DMEM high glucose media, 10% fetal calf serum (FCS), 1% penicillin and streptomycin (PS), 1% Glutamine. First passage cells were used in the experiments.

### Peptide and Protein Synthesis

The TAT protein transduction domain (RKKRRQRRR) was used to facilitate intracellular uptake of candidate molecules into primary human chondrocytes and mouse knee joints. For cell association studies, the TAT PTD and a control PTD were labeled with rhodamine and synthesized using solid-phase peptide synthesis with Fmoc protection in 25 μM scale on a Symphony Quartet peptide synthesizer ^14^. All peptides were cleaved and deprotected using a standard protocol (95% trifluoroacetic acid, 1% water, 1% triisopropylsilane). Crude peptides were precipitated using cold diethylether and purified using prep-scale reversed-phase HPLC with a C18 column; purity was confirmed by MALDI-TOF mass spectrometry using the matrix α-CHCA. Recombinant proteins were purified as described previously ^5,10.^ Plasmids containing either TAT-Cre ^31^ or the TAT PTD-DRBD (Protein-Transduction-Domain coupled to a Double-Stranded RNA Binding Domain) ^10^ were transformed into BL21 Codon Plus *e*. *coli*. and expanded to ∼1-2L scale. Expression was induced via IPTG (isopropyl β-D-1 thiogalactopyranoside), and cells were harvested after 2-3 hours. Multi-step purification proceed sequentially via affinity chromatography, cation exchange, and anion exchange yielding purified proteins for *in vivo* experiments ^10^. Purified proteins were quantified via SDS-PAGE electrophoresis and Coomassie staining against BSA standards. Activity was confirmed via *in vitro* assays in established cell lines.

### Cell Association Experiments

To assess the relevance of the TAT protein transduction domain as a drug delivery tool for synovial joints, fluorescently labeled TAT peptides and siRNAs were used in cell association experiments and assayed via flow cytometry and fluorescence microscopy as follows. 50,000 cells per well were plated in 12 well plates. Two days later cells were washed with HBSS and treated with control (GGSGGHHHHHHG) or TAT-peptides (1.25, 2.5, or 5 μM) for 1 hour or left untreated. Following treatment, cells were washed three times with HBSS containing 0.5 mg/mL heparin sulfate, detached with accutase, and resuspended in growth media for analysis. Cell-associated fluorescence was measured via flow cytometry. Live cells were discriminated via forward and side scatter; cell association was quantified using the mean fluorescence intensity. At each dosage, mean fluorescence values between control and TAT-peptides were compared using paired comparisons with an a priori significance level of 0.05. These studies were repeated for chondrocytes from n = 6 donors. To evaluate the dose-dependent association of fluorescently labeled TAT-peptides, primary human chondrocytes were cultured and treated with either a rhodamine labeled TAT peptide or labeled control peptide (1-10μM) for 10 minutes to 1 hour. The surface peptide was removed via HBSS washes and accutase digestion prior to analysis by flow cytometry.

### In vitro siRNA Knockdown

For a functional evaluation of the TAT PTD as a drug delivery method for synovial joints, siRNA knockdown experiments were performed using the TAT PTD-DRBD ^10^. siRNA was complexed with the PTD-DRBD at a 1:8 molar ratio, and cells were treated at concentrations of 50-400 nM siRNA. Control cells were either untreated or treated by cationic lipids (Lipofetamine 2000, Invitrogen) according to the manufacturer’s recommendations. For imaging, fluorescently-labeled siRNAs were used, and after 1 hour of treatment cells were washed with PBS and then trypsin, followed by immediate imaging. Otherwise, siRNAs targeting either GAPDH or FoxO1 were used. The off-target control siRNA was designed against GFP ^10^. siRNA effectiveness was determined via qPCR (GAPDH normalized to β-macroglobulin) or Western blotting (FoxO1). Functional siRNA-knockdown was examined via immunoblotting for FoxO1 and qPCR for GAPDH. All antibodies were from AbCam, and siRNAs were validated sequences from Ambion using previously described methodology ^2^.

### In vivo Genetic Recombination

To assess the *in vivo* ability of TAT PTD fusions, to modulate synovial joint biology, the protein TAT-Cre ^31^ was injected into the stifle joints of luciferase reporter mice ^26^. TAT-Cre was prepared as previously described. TAT-Cre was injected (10 μL of 60 μM) into the medial side of the left knees in the luciferase reporter mice (n=5). The contralateral knees were injected with buffer as a control. After treatment, *in vivo* luciferase imaging (IVIS Spectrum, Perkin Elmer, Waltham MA, USA) was performed daily for 4 days following intraperitoneal injection of D-luciferin (150 mg/kg). After imaging on day 5, a single mouse was euthanized and both knee joints were dissected to further evaluate the tissue source of the luciferase signal. The remaining mice were imaged monthly thereafter until 10 months following injection.

### Binding studies in Human OA Cartilage

Explants of Kellgren-Lawrence Grade IV OA cartilage were obtained from knee arthroplasty cases under IRB approval and equilibrated in sterile tissue culture.

Equilibrium peptide uptake was measured with previously published methods ^7^. Briefly, structurally intact 3 mm osteochondral cores (N=3) of human stage IV OA cartilage were taken from patients undergoing knee replacements. For control studies, samples were stored in phosphate buffered saline (PBS) with protease inhibitors for 24 hours prior to experimentation. To investigate the effect of the negative fixed charge density in cartilage, 500 mM NaCl was added to the PBS to disrupt the negative fixed charge of the proteoglycans. To investigate how bulk damage effects cartilage, samples were degraded in 5 mM trypsin for 24 hours prior to uptake studies.

During uptake studies, samples were incubated in PBS plus protease inhibitors and 100 mM labeled peptide, a control peptide or TAT, and graded amounts (150-400 mM) of unlabeled TAT for 48 hours. The control peptide is a peptide with the same fixed solubility and size as TAT but without the functionality of TAT. The control is referred to as control-TAMRA. The wet weight of each cartilage disk was measured. The fluorescence excitation of the solution was measured at 560 nm, and calibration curves were created based on known standards for both the TAT and control peptides. The sample was dried to measure the dry weight. The dried sample was then degraded with proteinase K to measure the uptake of peptide by fluorescence excitation. The uptake ratio was calculated, defined as the concentration of labeled peptide inside the cartilage divided by concentration of labeled peptide in the solution. We used linear regression to assess the dependence of uptake ratio on peptide concentration.

### Statistical Analysis

Analysis of variance (ANOVA) was used to detect statistical differences in quantitative measures between experimental groups. Planned comparisons between groups of interest were made using an *a priori* significance level of α = 0.05 with the Bonferroni correction for multiple comparisons. Cell association experiments were analyzed by performing a 2-factor ANOVA using the dependent variable of the mean live-cell fluorescence in the rhodamine channel via flow cytometry. Independent variables were peptide (TAT-PTD or control) and dosage, and paired comparison s were made between TAT-PTD and control at each dosage. Results are plotted as normalized to the average value achieved in the maximum group. GAPDH knockdown was assessed using a 1-way ANOVA with groups of untreated, control-lipofectamine, siGAPDH-lipofectamine, control-PTD-DRBD, and siGAPDH-PTD-DRBD. Bonferroni post-hoc tests were used to infer specificity of knockdown comparting the siGAPDH groups to the untreated control groups with an *a priori* significance level of α = 0.05. All data are expressed as mean ± standard error of the mean. *In vivo* bioluminescence imaging experiments were assessed by comparing the quantitative luminescence data for the buffer and TAT-Cre injected knees over time using a 2-way ANOVA with planned comparisons between the buffer and TAT-Cre injected knees at each timepoint. We used linear regression to assess the dependence of uptake ratio on peptide concentration.

## Results

### TAT-PTD Associates with Primary Human Chondrocytes

As a first test of the TAT-PTD, fluorescent dye TAT conjugates were incubated with primary chondrocytes *in vitro* followed by trypsinization, re-attachment and culture for 24 hours, and then either flow cytometry or fluorescent imaging. The TAT-PTD dye resulted in significantly more cell-associated fluorescence than the control peptide in a dose-dependent manner (Fig. 1A). The mean fluorescence intensity (MFI) for TAT-treated cells was significantly greater than for the chondrocytes treated with the control peptide at all doses (1.25-5 μM, all p ≤ 0.02). Cell association of TAT peptides was observed in both male and female chondrocytes from donors ranging in age from 24 to 64, and from normal and OA donors (Fig. 2). Because the cells were trypsinized prior to analysis, these data indicate that the TAT-PTD has strong potential for intracellular delivery of both experimental and therapeutic molecules in primary human chondrocytes.

**Figure 1.**
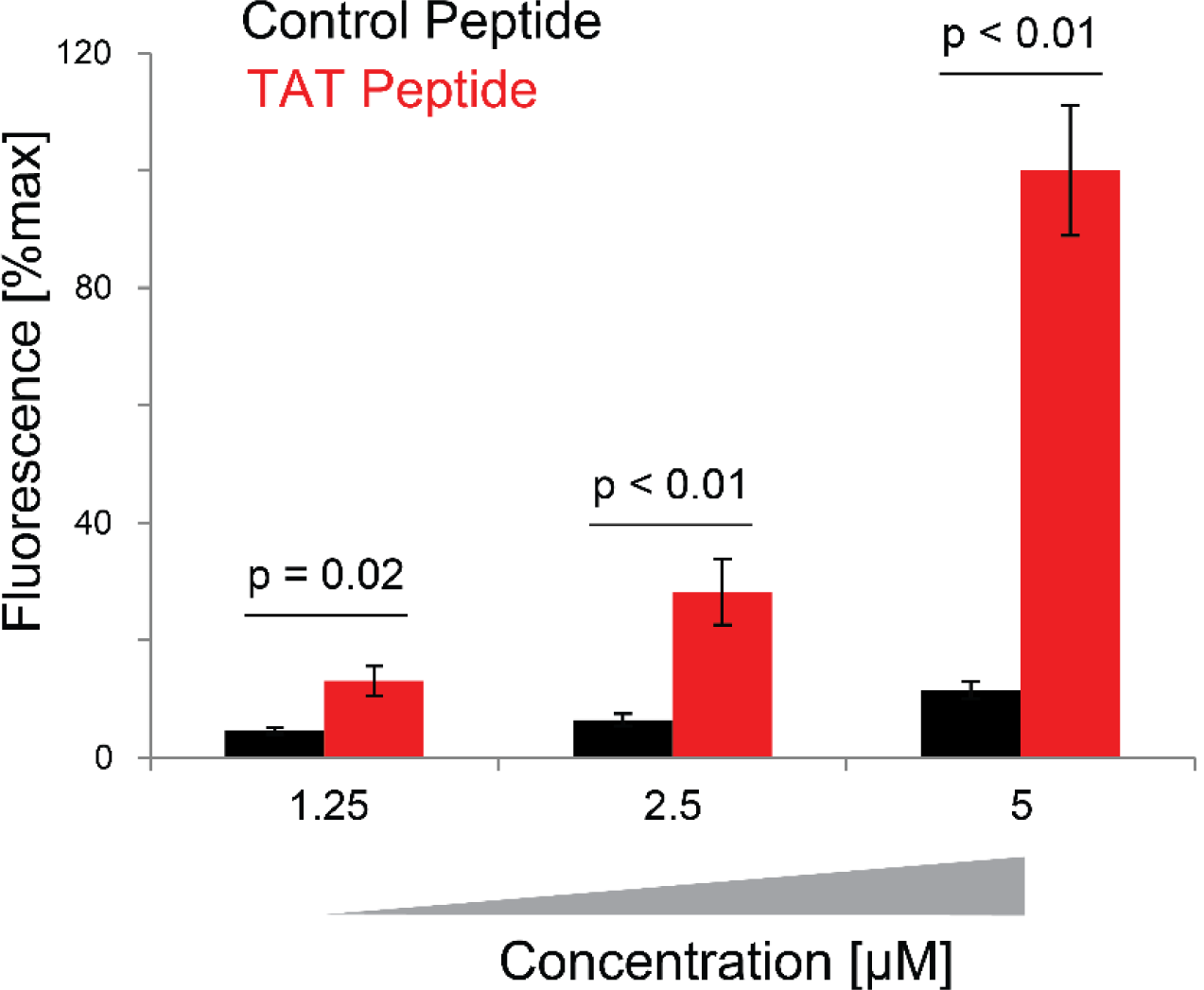
Dose-dependent association of TAT-PTDs with primary human chondrocytes. Primary human chondrocytes were cultured and treated with either a rhodamine labeled TAT peptide or labeled control peptide. Surface peptide was removed via HBSS washes and accutase digestion prior to flow cytometry. Significantly more fluorescence was found for all treatment dosages of TAT peptide compared to control peptide. Data from n=11 independent donors.

**Figure 2.**
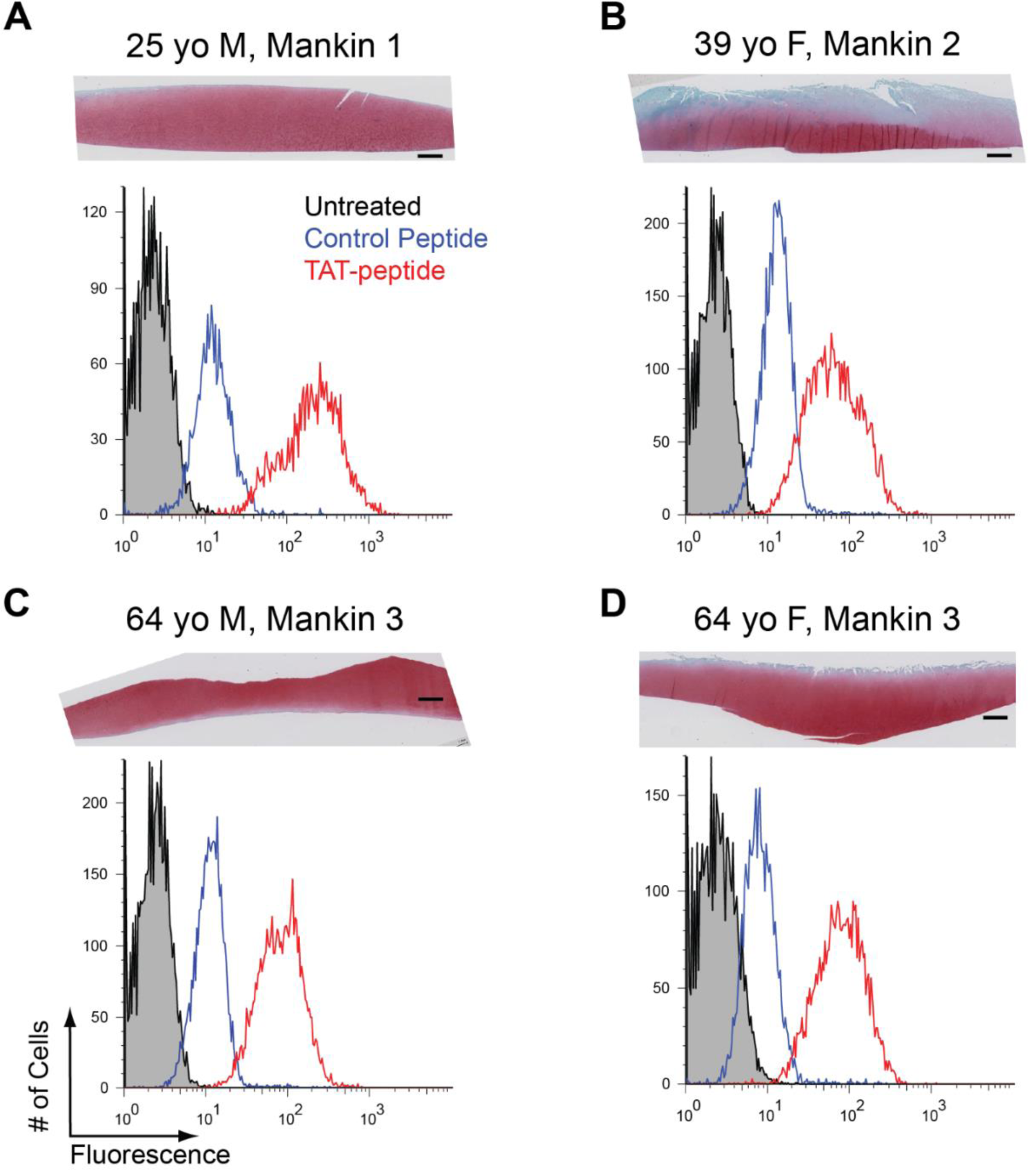
Cell association of TAT-PTDs in chondrocytes from a diverse array of donors. Each panel shows a Safranin-O Fast Green stained section (top) and flow cytometry histogram (bottom) from representative donors. For histograms, black population is untreated cells, blue is cells treated with a control rhodamine-labeled peptide, and red is a population treated with 5 μM rhodamine-labeled TAT peptide. (A) 25-year old male with Mankin 1 deterioration. (B) 39-year old female with Mankin 2 deterioration. (C) 64-year old male with Mankin 3 deterioration. (D) 64-year old female with Mankin 3 deterioration. Cell association experiments performed on n=10 donors.

### Functional Delivery of siRNA using the TAT-DRBD

To deliver siRNAs into monolayers of primary human chondrocytes in monolayer, the siRNA was bound by a TAT-PTD-DRBD (Double Stranded RNAi Binding Domain)^10^. The TAT-PTD was able to efficiently deliver fluorescently-labeled siRNA to primary human chondrocytes in monolayer. Fluorescence imaging (Fig. 3A) found diffuse fluorescence using the PTD-DRBD whereas Lipofectamine resulted in punctuate staining. Significant GAPDH mRNA knockdown was observed with both Lipofectamine (p=0.008) and the PTD-DRBD (p<0.0001, Fig. 3B). In addition, protein-level knockdown of FoxO1 was achieved with both delivery methods and resulted in concomitant decreases in GAPDH (Fig. 3C). These data indicate that the TAT-fusion protein PTD-DRBD is capable of delivering functional siRNAs for gene knockdown in primary articular chondrocytes.

**Figure 3.**
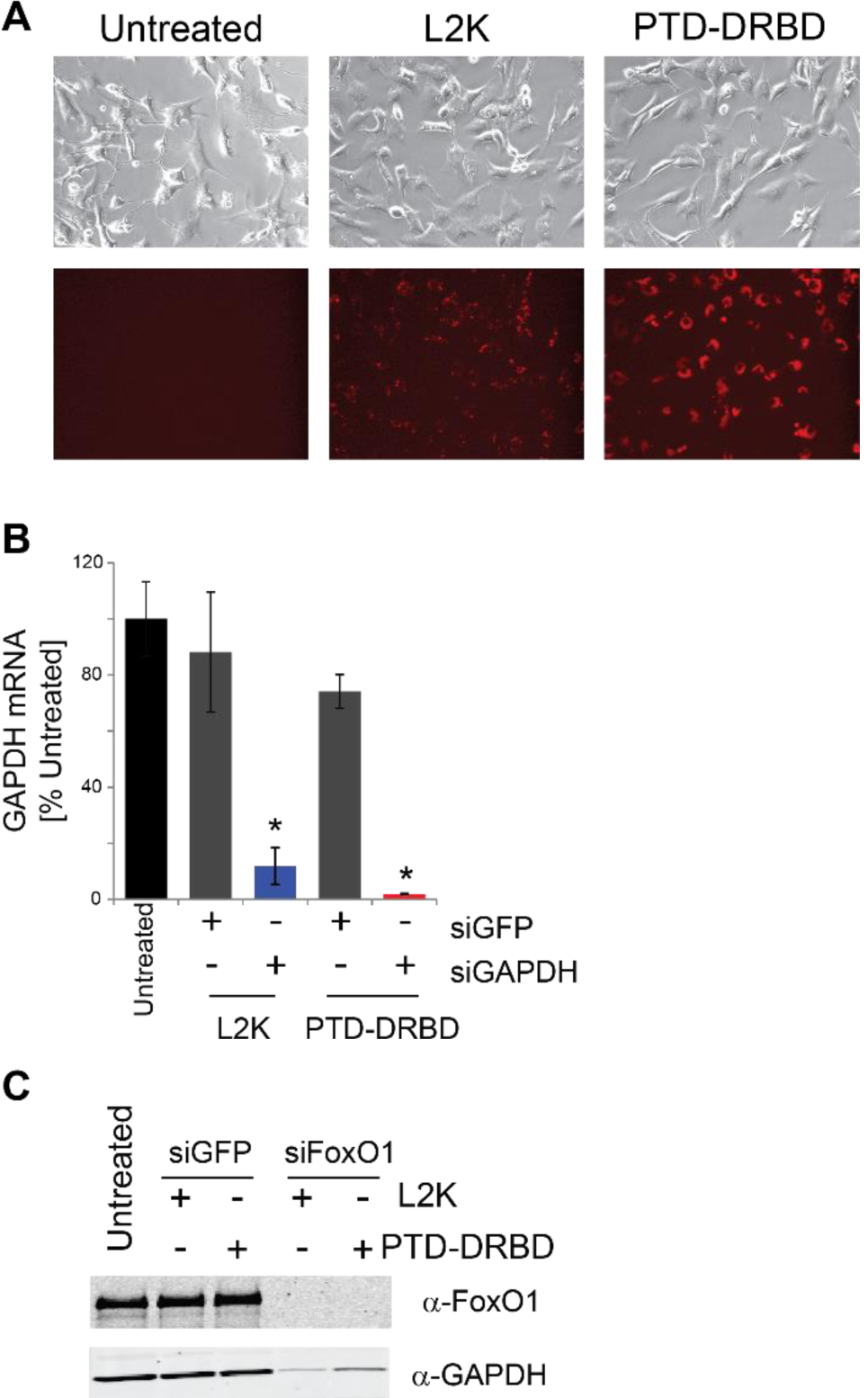
The TAT fusion protein PTD-DRBD delivers siRNA for RNA silencing in primary human chondrocytes. (A) siRNA was complexed to the PTD-DRBD TAT delivery protein (PTD- DRBD) or lipofectamine (L2K), and cells were treated as in Fig. 1A. Following PBS and trypsin washes, cells were imaged via phase contrast and epifluorescence microscopy. (B) Knockdown of GAPDH mRNA. Primary human chondrocytes were treated for 4 hours with siRNA designed against either GFP as a control (siGFP) or GAPDH (siGAPDH) as a target gene. mRNA levels were quantified by qPCR. Both lipofectamine and the PTD-DRBD resulted in significant knockdown of GAPDH mRNA. (C) Protein-level knockdown of FoxO1 measured by immunoblot. Primary human chondrocytes treated as in A with control siRNA (siGFP) or siRNA designed against FoxO1. Immunoblots performed following lysis and SDS-PAGE after 48 hours. Knockdown experiments using the TAT PTD-DRBD was performed on chondrocytes from n=10 donors with an siRNA concentration of 400 nM. Legend: L2K = lipofectamine 2000. PTD-DRBD = protein transduction domain coupled to a double-stranded RNA binding domain.

### *In Vivo* Efficacy of TAT-Cre using Reporter Mice

To test the *in vivo* efficacy of the TAT PTD, the recombinant enzyme TAT-Cre was injected intra-articularly into bioluminescent luciferase reporter mice. TAT-Cre injection resulted in significantly greater bioluminescence (p < 0.001) than buffer at all timepoints (all p < 0.01). Note that previous studies established that Cre-only does not enter cells ^31^.There was no main effect of time after injection on induced bioluminescence (p = 0.112). Bioluminescence was observed as rapidly as day 1 after injection and was sustained for 196 days (Fig. 4). Bioluminescence was observed in both the femoral condyles and the tibial plateau after injection of TAT-Cre.

**Figure 4.**
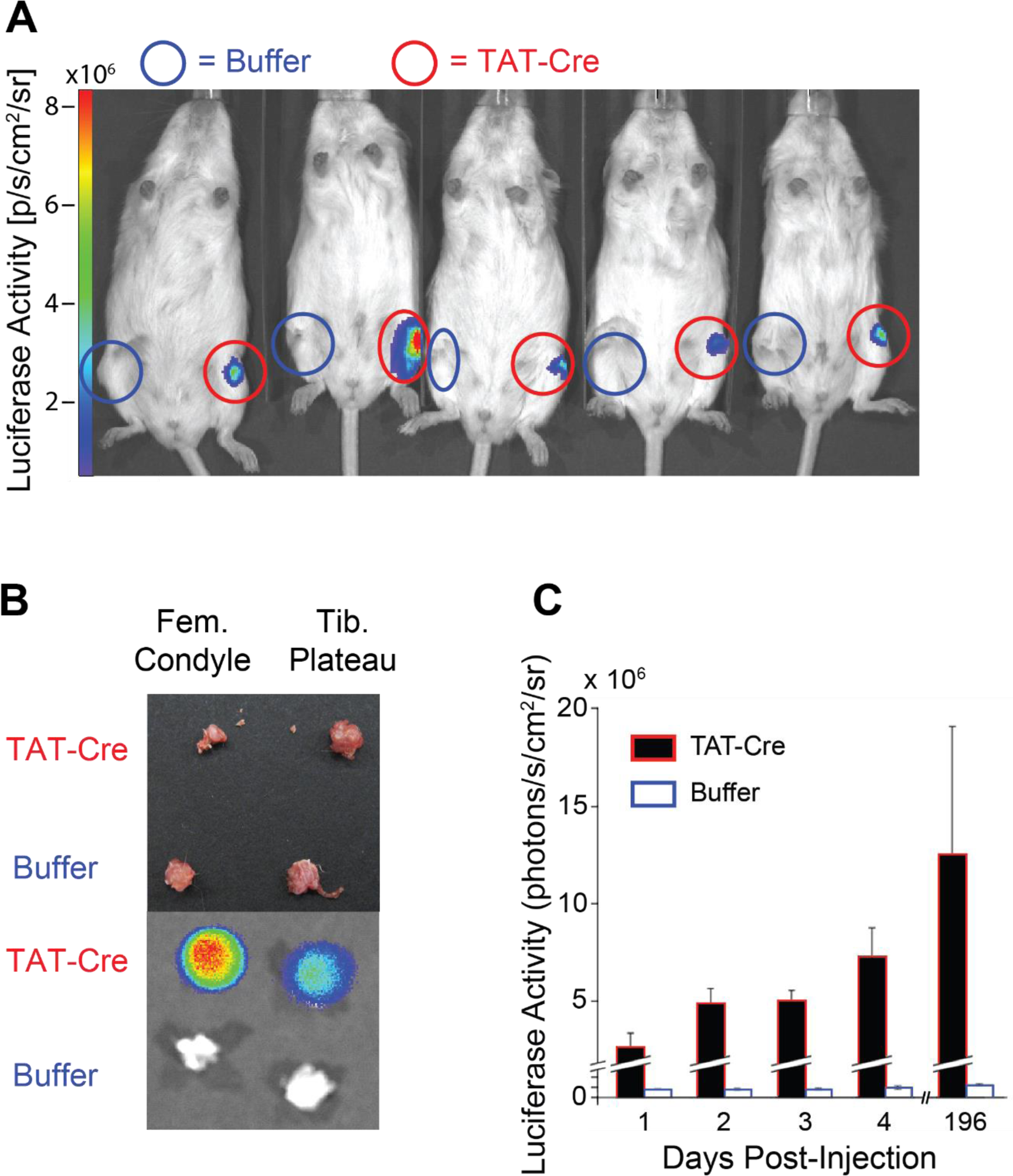
TAT-Cre induces genetic recombination in mouse knees. Cre-reporter mice were injected with either TAT-Cre or buffer in the intra-articular space prior to *in vivo* bioluminescence imaging. (A) Image of n=5 mice 4 days following injection. (B) Representative dissected mouse knee. Top shows photograph. Bottom shows bioluminescent image. (C) Quantification of TAT-Cre induced bioluminescence. Luciferase activity was quantified within circular regions of interest overlaid on both the control- and TAT-Cre- injected knees.

### Human OA Cartilage Preferentially Uptakes the TAT-PTD

To assess the transport properties of the TAT-PTD within human articular cartilage, fluorescent TAT and control peptides were incubated with cartilage explants (Fig. 5). The uptake ratio was not dependent on concentration of unlabeled TAT (slope: - 0.0006, 95% CI for uptake ratio: 1.67-3.11. n=48) or control peptide (slope: 0.0002, 95% CI: 0.004-2.31. n=48) (Fig. 5A). Therefore, the uptake ratio was equivalent to the partitioning coefficient ^7^. The mean partitioning coefficient of the labeled TAT peptide was 2.39, and the 95% confidence interval did not include 1, indicating that TAT causes uptake within human cartilage (Fig. 5A). The partitioning coefficient of TAT-PTD was reduced when the cartilage disk was treated with trypsin (p=0.045, n=24, Figure 5), but not 500 mM NaCl (p=0.3, n=24).

**Figure 5.**
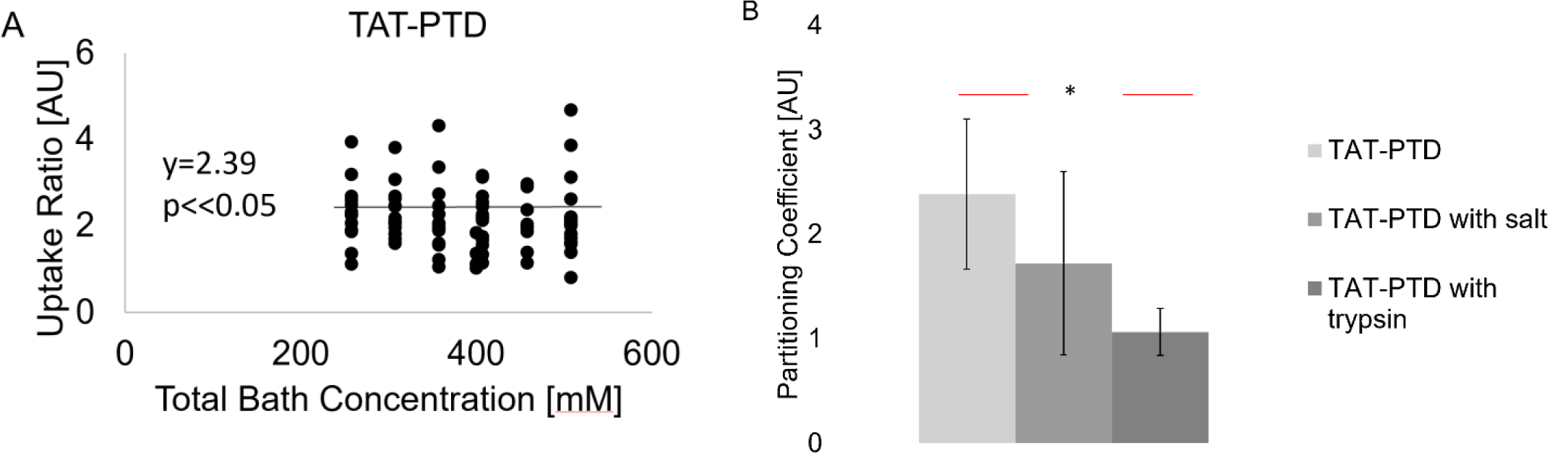
Transport studies of the TAT-PTD in human osteoarthritic articular cartilage. (A) Equilibrium uptake ratio between cartilage samples and bath for the TAT-PTD across a range of peptide concentrations in the bath. The uptake ratio of TAT-PTD within cartilage showed a slope < 0.01, indicating that partitioning is not concentration dependent. Therefore, the partitioning coefficient is the uptake ratio. (B) Both high-salt and trypsin resulted in decreased partitioning coefficients compared to untreated cartilage. This suggests that electrostatic interactions between cartilage GAGs and the TAT-PTD mediate the uptake of TAT-PTDs into cartilage.

## Discussion

The objectives of this study were first to determine if the TAT-PTD could transduce macromolecules into chondrocytes *in vitro*, and second to assess the ability of the TAT-PTD to bind to cartilage and transduce cells that are embedded in the cartilage extracellular matrix. The *in vitro* studies show that the TAT-PTD associated with primary human chondrocytes and is capable of delivering functional siRNA for knockdown of gene and protein expression by RNA interference. Using luciferase reporter mice, *in vivo* injection of TAT-Cre resulted in induction of luciferase activity indicating that the TAT-PTD can deliver functional enzymes directly to synovial tissues. Binding studies in human osteoarthritic cartilage showed that the TAT-PTD preferentially binds to cartilage in a trypsin-dependent manner, suggesting that the TAT-PTD may be capable of extending the residence time of molecules injected into the joint.

The binding studies in human OA cartilage retrieved during joint replacement found preferential binding of the TAT-PTD. To quantify uptake into cartilage, we defined the uptake ratio as the ratio of the concentration of labeled peptide inside the cartilage to the concentration of labeled peptide in the bath. Because the lower bound of the 95% confidence interval for the uptake ratio of the TAT-PTD is greater than one, these data show that the TAT-PTD increased uptake and partitioning of a fluorescent marker TAMRA. Furthermore, the equilibrium distribution of TAT-PTD between cartilage and the fluid indicated a partitioning coefficient greater than 1. Additionally, trypsin treatment decreased uptake of TAT-PTD. This is notable because this is a common model for disease conditions such as OA. Because these studies were performed in late-stage OA cartilage, it was likely that the TAT-PTD may have even greater uptake and partitioning in healthy and early-stage cartilage, although future studies are needed to confirm this.

These studies have important limitations. First, while the binding studies were performed in PBS, it is unclear if the TAT-PTD will have similar transport properties within synovial fluid of the joint. Thus, there is the possibility that delivery using the TAT-PTD may be less efficient *in vivo*. The cell-association measurements cannot conclusively be interpreted as evidence of chondrocyte TAT-PTD internalization. However, because association measurements were performed after trypsinization these data, as well as the siRNA knockdown studies, strongly suggest that the TAT-PTD is internalized by primary human chondrocytes. While the positive results from the injection of TAT-Cre into reporter mice suggest that *in vivo* delivery is possible, this study did not determine which joint cells were transduced, and the efficacy of the TAT-PTD to deliver in human joints remains unknown. Furthermore, although transduction was demonstrated in both male and female chondrocytes, it is unclear if there are differences between male and female joint cells and tissues. A final limitation of these studies is that all studies were examined at equilibrium. The specific dynamics of TAT-PTD binding to articular cartilage remain unknown. Addressing these limitations in future studies will enable more-precise modeling and prediction of the dynamics of the TAT-PTD within synovial joints, and this information will be valuable for assessing residence times of potential therapeutic compounds.

Previous work with AVIDIN, a small positively charged biotin binding protein, shows that electrostatic interactions enhance uptake in intra-articular injections with higher uptake of charged particles in the negatively charged cartilage as compared to surrounding tissues ^3^. Similarly, the TAT-PTD is positively charged and has the advantage of also inducing intracellular uptake of its bound molecules ^27^. Future work will test the correlation between GAG content and TAT-PTD uptake with DMMB blue ^7^.

The data demonstrate that TAT-PTD can be used to deliver siRNA and protein to modulate chondrocyte biology. Consistent knockdown of GAPDH was achieved using the PTD-DRBD. TAT fusions penetrate articular cartilage and display prolonged joint residence time. The TAT protein transduction domain facilitates peptide-uptake in primary human chondrocytes from male and female donors with varying ages and OA grades. Taken together, our data suggest that cytosolic delivery using the TAT-PTD has strong potential for both basic and translational studies of synovial joint biology and osteoarthritis.

